# Activation of LH GABAergic inputs counteracts fasting-induced changes in tVTA/RMTG neurons

**DOI:** 10.1101/2021.11.24.469955

**Authors:** Nathan Godfrey, Min Qiao, Stephanie L. Borgland

**Affiliations:** University of Calgary, Department of Physiology and Pharmacology, Calgary, Alberta, T2N 4N1

**Keywords:** ventral tegmental area, fasting, dopamine, inhibitory synaptic transmission, tVTA, RMTg, GABA, motivational state

## Abstract

Dopamine neurons in the ventral tegmental area (VTA) are strongly innervated by GABAergic neurons in the ‘tail of the VTA’ (tVTA), also known as the rostralmedial tegmental nucleus (RMTg). Disinhibition of dopamine neurons through firing of the GABAergic neurons projecting from the lateral hypothalamus (LH) leads to reward seeking and consumption through dopamine release in the nucleus accumbens. VTA dopamine neurons respond to changes in motivational state, yet less is known of whether tVTA/RMTg GABAergic neurons or the LH GABAergic neurons that project to them are also affected by changes in motivational state, such as fasting. An acute 16 h overnight fast decreased the excitability of tVTA/RMTg GABAergic neurons of male and female mice. In addition, fasting decreased synaptic strength at LH GABA to tVTA/RMTg GABAergic synapses, indicated by reduced amplitude of optically evoked currents, decreased readily releasable pool (RRP) size and replenishment. Optical stimulation of LH GABA terminals suppressed evoked action potentials of tVTA/RMTg GABAergic neurons in unfasted mice, but this effect decreased following fasting. Furthermore, during fasting, LH GABA inputs to tVTA/RMTg neurons maintained functional connectivity during depolarization, as depolarization block was reduced following fasting. Taken together, inhibitory synaptic transmission from LH GABA inputs onto tVTA/RMTg GABAergic neurons decreases following fasting, however ability to functionally inhibit tVTA/RMTg GABAergic neurons is preserved, allowing for possible disinhibition of dopamine neurons and subsequent foraging.

**Key Points:**

- While dopamine neuronal activity changes with motivational state, it is unknown if fasting influences tVTA/RMTg GABAergic neurons, a major inhibitory input to VTA dopamine neurons.
- In unfasted mice, there were sex differences in inhibitory synaptic transmission onto tVTA/RMTg GABAergic neurons.
- Activation of LH GABAergic neurons decreases firing of tVTA/RMTg GABAergic neurons through a monosynaptic input.
- An acute fast decreased the excitability of tVTA/RMTg GABAergic neurons.
- An acute fast decreases inhibitory synaptic transmission of the LH GABA input to tVTA/RMTg GABAergic neurons in both male and female mice.

## Introduction

Motivated behaviors arise during physiological need. The ventral tegmental area (VTA) dopamine neurons play a critical role in encoding the salience of cues in the environment that predict the quality and availability of rewards and dopamine release in the nucleus accumbens (NAc) can motivate animals to seek out these rewards (Salamone *et al*., 2003; Wise, 2006; Berridge, 2009). Dopamine neurons alter their firing activity and output in a state-dependent manner. For example, negative sodium balance increases phasic dopamine release in the NAc, whereas dopamine concentration was decreased when rats were sodium replete, suggesting that dopamine neurons track fluid balance in a state-dependent manner (Cone *et al*., 2016; Fortin & Roitman, 2018). Similarly, VTA dopamine neurons respond to a motivational state induced by food deprivation. During an acute fast, there is a sex dependent increase in excitatory synaptic transmission onto dopamine neurons (Godfrey & Borgland, 2020) as well as increased somatodendritic dopamine release measured by its effects on D2 receptor mediated inhibitory post synaptic currents (IPSCs) (Roseberry, 2015). Optogenetic activation of VTA dopamine neurons increases food retrieval in fasted animals, but prior acute exposure to high fat diet strongly suppresses this effect (Mazzone *et al*., 2020). Taken together, VTA dopamine neuronal activity changes with motivational state.

The VTA is composed of approximately 60-70% dopaminergic neurons, 25-30% GABAergic neurons and 2-5% glutamatergic neurons (Margolis *et al*., 2006, 2012; Nair-Roberts *et al*., 2008; Chieng *et al*., 2011; Ungless & Grace, 2012). The caudal region or the ‘tail of the VTA’ (tVTA) also known as the rostralmedial tegmental nucleus (RMTg) is composed of almost exclusively GABAergic neurons (Jhou *et al*., 2009). In brain atlases, the tVTA/RMTg is not differentiated from the VTA itself and thus non-projecting GABAergic neurons within the caudal region of the VTA are likely part of this structure. Dopamine neurons are the principal target of tVTA/RMTg neurons (Balcita-Pedicino *et al*., 2011). As such, these GABAergic neurons exert a significant inhibitory drive onto VTA dopaminergic neurons (Barrot *et al*., 2012). Optogenetic stimulation of VTA GABAergic neurons disrupts licking for sucrose solution in fasted mice (Van Zessen *et al*., 2012). Furthermore, a variety of aversive stimuli activate tVTA/RMTg neurons (Sánchez-Catalán *et al*., 2017; Li *et al*., 2019c). Given that hunger induced by acute fasting is an aversive motivational state (Betley *et al*., 2015), it is unknown if GABAergic neurons of the tVTA/RMTg are also sensitive to state-dependent changes in activity from fasting.

LH GABAergic neurons send strong projections to VTA GABAergic neurons and likely to the tVTA/RMTg, as these regions are in close proximity and often not distinguished (Kaufling *et al*., 2009; Nieh *et al*., 2015; Barbano *et al*., 2016; Marino *et al*., 2020). Activation of LH GABAergic terminals in the VTA disinhibits VTA dopamine neurons and increases dopamine release in the NAc (Stuber & Wise, 2016). Electrical or optogenetic stimulation of LH GABAergic neurons elicits robust feeding behaviour (Nieh *et al*., 2015; Stuber & Wise, 2016). Some reports have indicated that activation of the LH GABAergic to VTA GABAergic projection elicits feeding behaviour through disinhibition of dopamine neurons (Nieh *et al*., 2015, 2016), while others suggest that activation of LH GABAergic neurons mediates feeding behaviour through an *en passant* projection through the VTA terminating near the locus coeruleus (Marino *et al*., 2020). Furthermore, while it was proposed that feeding evoked by stimulation of GABAergic LH to VTA projections was due to the rewarding properties of food rather than the aversive drive state associated with hunger, it was never specifically tested whether this input was sensitive to an acute fast (Nieh *et al*., 2016). Here, we tested whether caudal tVTA/RMTg GABAergic neurons were sensitive to an acute fast and how synaptic transmission of LH GABA to tVTA/RMTg GABAergic neuronal projection was altered by this motivationally relevant stimulus.

## Methods

### Ethical Approval

All experiments and procedures were in accordance with the ethical guidelines established by the Canadian Council for Animal Care and were approved by the University of Calgary Animal Care Committee (protocol no. AC17-0034) and are consistent with the Journal’s policies for the ethical treatment of animals (Grundy, 2015).

### Subjects

Male and female mice 2-4 months old were housed in groups of two to five in same sex cages and were maintained on a 12 h light-dark schedule (lights on at 08.00 h, zeitgeber time (ZT) 0 and were given chow and water ad libitum. Mice were fed chow (5062 from Pico-Vac lab diet, Lab Supply, Fort Worth Tx), which is composed of (% of total kcal) 23% protein, 22% fat (ether extract), and 55% carbohydrate. The total caloric density of this diet was 4.60 kcal g^−1^. Experiments were performed during the animal’s light cycle. Male and female VGAT^cre^TdTomato mice, used to identify GABA neurons, were generated by crossing VGAT^cre^ (Slc32a1tm2(cre)Lowl/J) obtained from Jackson Laboratory (strain: 016962) with Rosa-td Tomato mice (B6.Cg-Gt[ROSA]26 Sortm9[CAG-tdTomato]Hze/J [Ai9] (strain: #007909). All mice were bred locally in the Clara Christie Centre for Mouse Genomics. Mice were anesthetized with isoflurane (vapourized at 3-5%, with O_2_ at 0.5-1 L/min), injected with subcutaneous meloxicam (5 mg/kg) and saline (2-3 mL, 0.9%), and secured in the stereotaxic frame (Kopf; Tujunga, CA). Channelrhodopsin (ChR2) (AAV2-EF1a-DIO-hChR2(H134)-EYFP; Neurophotonics, Centre de Recherche CERVO, Quebec City, QC, Canada) was bilaterally infused into the lateral hypothalamus (LH) with the following coordinates: ML: +/-1.000 mm, AP: −0.800 mm, DV: - 5.300 mm. Mice were returned to home cages for 6 to 7 weeks for full virus expression before use in experiments. Mice were initially weighed and then isolated 10 h into their light phase (ZT10), and single-housed overnight for a total of 16 h. During this time, food was removed from the cages of the fasted group and chow was given ad libitum to the control group. Water was given ad libitum to both groups. Water intake and body weight were recorded for fasted and control groups and food consumption per body weight was recorded in the control group after the 16 h fast (ZT2). Electrophysiology experiments were performed on mice at ZT2.

### Electrophysiology

All electrophysiological recordings were performed in slice preparations from adult (2-4 months old) VGAT^cre^Td-Tomato mice. In these mice, a red fluorescent marker is expressed exclusively in GABA neurons, permitting immediate identification of GABAergic neurons in the tVTA/RMTg for electrophysiological recordings. We recorded fluorescently identified GABAergic neurons from the caudal/medial VTA located medial to the medial lemniscus (ML). Mice were deeply anaesthetized with isoflurane and transcardially perfused with an ice-cold N-methyl-D-glucamine (NMDG) solution containing (in mM): 93 NMDG, 2.5 KCl, 1.2 NaH_2_PO_4_.2H_2_O, 30 NaHCO_3_, 20 Hepes, 25 D-glucose, 5 sodium ascorbate, 3 sodium pyruvate, 2 thiourea, 10 MgSO_4_.7H_2_O, 0.5 CaCl_2_.2H_2_O and saturated with 95% O_2_-5% CO_2_. Mice were then decapitated, and the brain extracted. Horizontal midbrain sections (250 μm) containing the VTA were cut on a vibrating-blade microtome (Leica, Nussloch, Germany). Slices were then incubated in NMDG solution (32°C) and saturated with 95% O_2_-5% CO_2_ for 10 min. Following this, the slices were transferred to artificial cerebrospinal fluid (aCSF) containing (in mM): 126 NaCl, 1.6 KCl, 1.1 NaH_2_PO_4_, 1.4 MgCl_2_, 2.4 CaCl_2_, 26 NaHCO_3_, 11 glucose, (32°C) and saturated with 95% O_2_-5% CO_2_. Slices were incubated for a minimum of 45 min before being transferred to a recording chamber and superfused (2 mL min^−1^) with aCSF (32-34°C) and saturated with 95% O_2_-5% CO_2_. Neurons were visualized on an upright microscope using ‘Dodt-type’ gradient contrast infrared optics and whole-cell recordings were made using a MultiClamp 700B amplifier (Molecular Devices, San Jose, CA, USA). The recorded signal was collected at a sampling rate of 20 kHz and a 2 kHz Bessel filter was applied to the data during collection. For optically evoked currents, a 3 ms, 3.5 mW, pulse of 470 nm was provided by an LED light source through a ×40/0.80 water immersible Olympus microscope objective. Recorded neurons were positioned so that the soma was at the centre of the objective. All electrophysiology experiments were done in the presence of DNQX (10 μM).

#### Voltage clamp recordings

For all voltage clamp recordings, recording electrodes (2-5 MΩ) were filled with Cesium Chloride internal solution containing (in mM): 140 CsCl, 10 HEPES, 0.2 EGTA, 1 MgCl_2_, 2 MgATP, 0.3 NaGTP, 5 QX-314-Cl, and 0.2% Biocytin. The liquid junction potential was 3.5 mV, but not corrected for.

#### Validation of ChR2 expression and function

To validate and confirm the selective expression and function of ChR2 in GABA neurons of the LH, train stimulations were evoked at 5 Hz, 10 Hz, and 20 Hz recording from LH GABA neurons or tVTA/RMTg GABA neurons. Example traces are averages of 3 sweeps. A 1 second, continuous, stimulation was also optically evoked in both LH GABA neurons and tVTA/RMTg GABA neurons. Furthermore, optically evoked currents were recorded before and after the bath application of picrotoxin (100 μM). The two minutes, 12 sweeps, prior to picrotoxin application were compared to the two minutes, 12 sweeps, after the effect of picrotoxin was seen to take full effect. Finally, optically evoked currents were recorded in the presence of TTX (1 μM) followed by TTX and 4-AP (100 μM) to determine monosynaptic connection between the LH GABA neurons and the tVTA/RMTg GABA neurons.

#### Miniature inhibitory (mIPSC) postsynaptic currents

Recording electrodes (2-5 MΩ) were filled with a CsCl internal solution. Neurons were voltage clamped at −70 mV and following the application of tetrodotoxin (TTX; 1 μM), the current was recorded for 5 min. mIPSCs were identified and measured using Mini Analysis 60 (Synaptosoft, Decateur, GA, USA) and the following parameters: amplitude > 10 pA, decay time < 10 ms, and rise time < 4 ms.

#### Strontium experiments

Recording electrodes were filled with CsCl internal solution. The 2.4 mM CaCl_2_ in the aCSF was replaced with 2.4 mM SrCl_2_ when slices were transferred from the NMDG solution. Neurons were voltage clamped at −70 mV and currents were evoked with a single optical stimulation at 0.03 Hz (3.5 mW) in the presence of 4-AP. To ensure that neurons analyzed received inputs with adequate ChR2 expression, neurons with a current amplitude less than 30 pA immediately following optical stimulation were excluded. Using Mini Analysis 60 and the following parameters: amplitude > 10 pA, decay time < 10 ms, and rise time < 4 ms, quantal currents were identified for the first 1000 ms following the optical stimulation. The amplitude and frequency of Sr^2+^ events were compared between early (50 to 350 ms) and late (700 to 1000 ms) periods of asynchronous activity following optical stimulation.

#### Optical stimulation trains

To evoke oIPSCs, recording electrodes (2-5 MΩ) were filled with a CsCl internal solution. Neurons were voltage clamped at −70 mV and a 20 Hz over 1 s (3.5 mW) optical stimulation at 0.1 Hz was used to evoke IPSC trains. A 1 minute, 6 sweep, period was used to create average traces with MatLab (https://github.com/borglandlab/Godfrey-Qiao-and-Borgland-2021.git) and current peaks were identified. To ensure that neurons analyzed received inputs with adequate ChR2 expression, neurons with an average first peak amplitude less than 50 pA were excluded. Peak amplitude of the first oIPSC, the readily releasable pool (RRP) and the steady-state oIPSC amplitude were calculated using methods established previously (Thanawala & Regehr, 2013, 2016). For the RRP_train_, averaged oIPSC amplitudes were summed throughout the train stimulus to give a cumulative oIPSC curve. A line was fit to the final 6 points of the cumulative oIPSC and back extrapolated to the y-axis. The slope of this line is the steady-state oIPSC amplitude, reflecting the rate of vesicle recycling after the RRP has been depleted, and the y-intercept is the RRP_train_ amplitude. Two neurons were excluded from the RRP analysis as the calculation of the y-intercept produced a negative y-intercept, which is physiologically not possible. The paired pulse ratio for the first 5 optical stimulations of the train was calculated as pulse (P) n / pulse (P) 1 (McGarry & Carter, 2016, 2017).

#### Current Clamp Recordings

For all current clamp recordings, recording electrodes (2-5 MΩ) were filled with Potassium-gluconate internal solution containing (in mM): 130 K-gluconate, 10 KCl, 10 HEPES, 0.5 EGTA, 10 Na_2_-Phosphocreatine, 4 MgATP, 0.3 NaGTP, and 0.2% Biocytin. The liquid junction potential was 15.9 mV. Recording electrodes (2-5 MΩ) were filled with a K-gluconate internal solution. Current was injected to hold the cell at −70 mV and 500 ms current steps, from −75 pA to 200 pA in 25 pA increments at 0.1 Hz, were used to evoke action potentials. For the −75 pA to 0 pA steps, the membrane potential at the end of each step was plotted against the current step. The input resistance was determined to be the slope of the line through these points. Action potentials were identified using Clampfit. Neurons that failed to fire action potentials by a 100 pA current step were deemed to be unhealthy and were excluded from analysis. To measure the resting membrane potential, the membrane potential of the cell was averaged over 5 minutes using Clampfit. Neurons with spontaneous action potentials were excluded from this calculation.

For optical stimulation during current clamp recordings, current was injected to hold the cell at −70 mV and 1 s current steps to 100 pA at 0.1 Hz were used to evoke action potentials. After allowing the cell to stabilize, a 1-minute (6 sweeps) baseline was recorded. Following the baseline, a 20 Hz optogenetic stimulation was synchronized with the onset of the current step and the membrane potential was recorded for 1 minute (6 sweeps). Using MatLab, action potentials were identified. The latency to fire (time to first peak), peak amplitude (difference between threshold and peak), and the after-hyperpolarization potential (AHP; difference between the inflection point of the AP on the falling phase and threshold voltage) were also calculated and averaged over 1s or the first 200 ms with or without optical stimulation. Neurons that did not have more than 1 action potential in the first 200 ms of the current step were excluded from this analysis. Depolarization block occurred in neurons that ceased firing in the final 200 ms of the current step. A two-tailed binomial test, Wilson/Brown, was used to compare between the proportion of depolarization block and no depolarization block with and without optogenetic stimulation.

### Immunohistochemistry

Mice were deeply anesthetized with isoflurane and transcardially perfused with phosphate buffered saline (PBS) and then with 4% paraformaldehyde (PFA). Brains were dissected and post-fixed in 4% PFA at 4°C overnight, then switched to 30% sucrose. Frozen sections were cut at 30 μm using a cryostat. 10% goat and donkey serums were applied to block non-specific binding for 1 hour. To examine colocalization of TdTomato in VGAT^cre^TdTomato mice with tyrosine hydroxylase (TH), horizontal sections were then incubated with primary antibody mouse anti TH 1:1000 (Sigma, T1299) and rabbit red fluorescent protein (RFP) 1:2000 (Rockland, 600-401-379) to amplify the tdTomato signal in 1% BSA for 24 hours at room temperature followed by incubation with secondary antibody Alexa Fluor 488 goat anti mouse 1:400, Alexa Fluor 594 donkey anti-rabbit 1:400, and DAPI 1:2000 for 1 hour. To check viral transfection, coronal sections were incubated with primary antibody chicken GFP 1:1000 (AVES Labs, GFP-1020) and rabbit RFP (1:2000) in 1% BSA for 1 hour at room temperature followed by incubation with secondary antibody Alexa Fluor 488 goat anti chicken 1:400 and Alexa Fluor 594 donkey anti-rabbit 1:400 for 1 hour. Slices were mounted with Fluroshield (Sigma). To check biocytin-filled neurons, slices were fixed in 4% PFA overnight at 4° C and then rinsed in PBS, incubated with Alexa Flour 488 streptavidin (1:200) for 2 hours at room temperature. All images were obtained on an Olympus virtual slide microscopy VS120-L100-W with a 10x objective (Olympus Canada Inc., Ontario, Canada) and a Leica confocal microscopy TCS SP8 with a 25x objective (Leica Microsystems Inc., Ontario, Canada). Images were processed using ImageJ/FIJI. Neurons in the entire 25 x image were then counted using the colocalization object counter ImageJ/FIJI plugin and following protocol established by Lunde and Glover (Lunde & Glover, 2020). Neurons were identified as VGAT+ by co-expression of td-Tomato and DAPI; TH and DAPI; and VGAT+ with TH+ by co-expression of td-Tomato, TH, and DAPI. To determine if TdTomato fluorescence and TH-like staining co-localized, we averaged cell counts from 63 slices from 6 mice, with an average of 21 neurons expressing td-Tomato and DAPI per slice.

Following the voltage clamp 20 Hz train stimulation experiments, the slices were immediately placed in 4% PFA to fix for a minimum of 24 hours at 4°C. The slices were then transferred to PBS and stored at 4°C. Prior to imaging, slices were rinsed 3 times in PBS and then mounted with Fluoshield (Sigma). Whole brain images were obtained on an Olympus virtual slide microscopy VS120-L100-W with a 10x objective (Olympus Canada Inc., Ontario, Canada) with identical parameters. Using ImageJ, a region of interest was created by tracing the outside of the slice and the value of the mean fluorescent intensity was obtained as an ImageJ measurement.

### Data Analysis

All values are expressed as means +/- SD and assessed for normality using a Shapiro-Wilk test. In Fig 1 symbols represent individual mice, while in Figs 2–9 symbols represent individual neurons. Outliers were identified and removed using the Robust regression and Outlier removal (ROUT) (Q = 1%) of current amplitudes and in the change in latency to fire. Statistical significance was assessed by using two-tailed unpaired Student’s t test for two comparisons. A two-way ANOVA followed by Sidak’s multiple comparisons was used for multiple group comparisons. For time course experiments, either a repeated measures two-way ANOVA followed by Tukey’s multiple comparisons test, a mixed effects analysis followed by Tukey’s multiple comparisons test, or a repeated measures one-way ANOVA followed by Dunnett’s multiple comparisons test were used. For comparing proportions, a two-tailed binomial test was used. In all electrophysiology experiments, sample size is expressed as N/n, where N refers to the number of neurons recorded from n animals. Asterisks were used to express statistical significance in figures: *P < 0.05, **P < 0.01, ***P< 0.001, and ****P < 0.0001. GraphPad Prism 8.3 (GraphPad Software, Inc., La Jolla, CA, USA) was used to perform statistical analysis. Figures were generated using GraphPad Prism 8.3 and Adobe Illustrator CS4 (Adobe Inc., San Jose, CA, USA) software.

**Figure 1.**
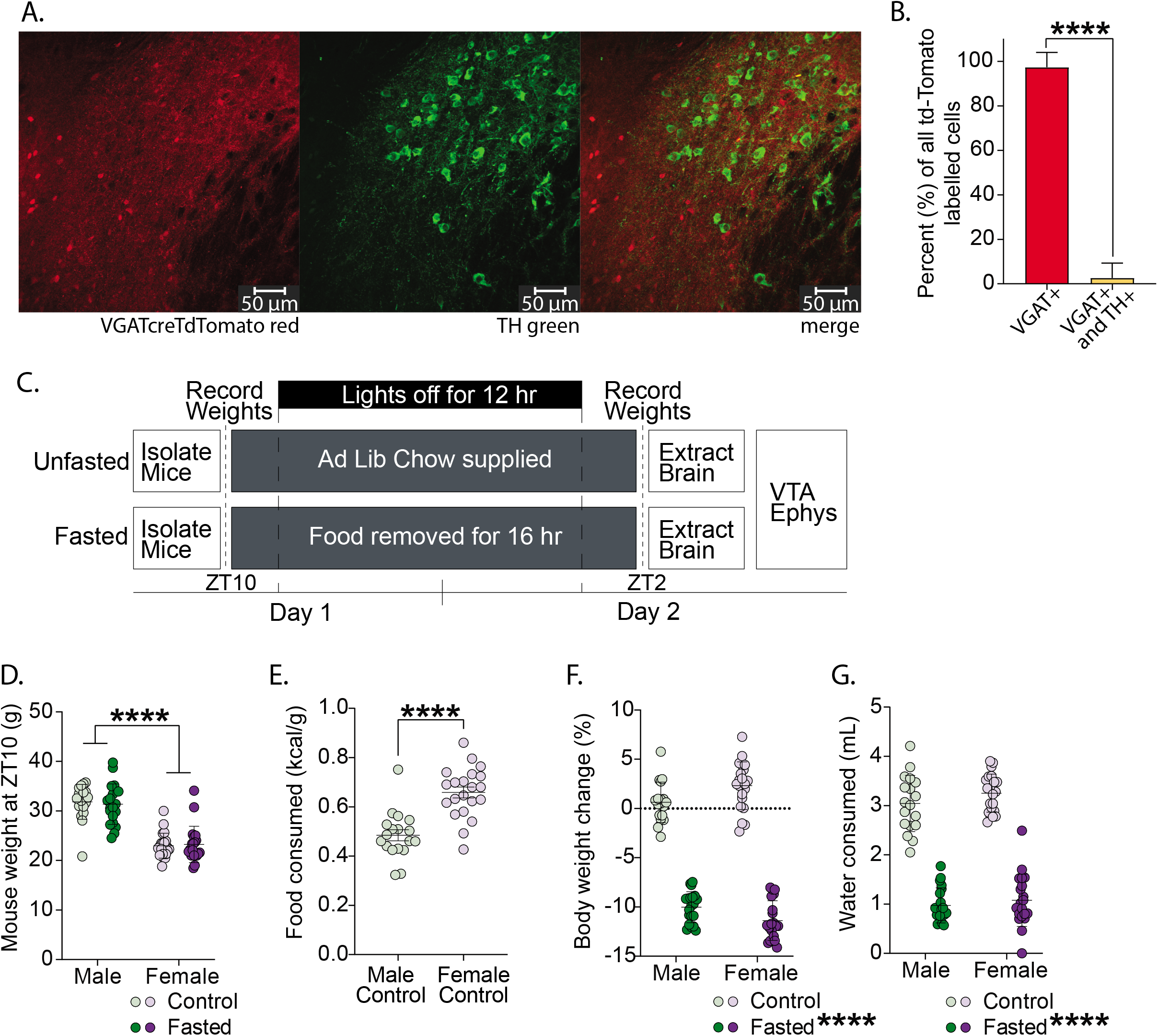
Validation of VGAT^cre^TdTomato mouse line and effects of an acute fast on body weight and consumption in male and female mice. **A**, Confocal images of cells expressing VGAT^cre^TdTomato (red), TH (green), and merged in the caudal VTA. **B**, percent of all td-Tomato labelled cells in the VTA that are only VGAT+, and VGAT+ and TH+. Only a very small proportion of cells are both VGAT+ and TH+. **C**, schematic representation of time course of fast and when weights were recorded. **D**, mouse weight (g) prior to the 16 h overnight isolation or fast. Male mice weigh more than female mice. **E**, chow consumed (kcal/body weight (g)) by control male and female mice during the 16 h overnight period of isolation. Female mice consume more food per body weight than male mice. **F**, body weight change between before and after the 16 h overnight period. Male and female fasted mice have reduced body weight after fasting. **G**, water consumed by male and female mice during the 16 h overnight period. Male and female fasted mice reduce water intake during the acute fast.

**Figure 2.**
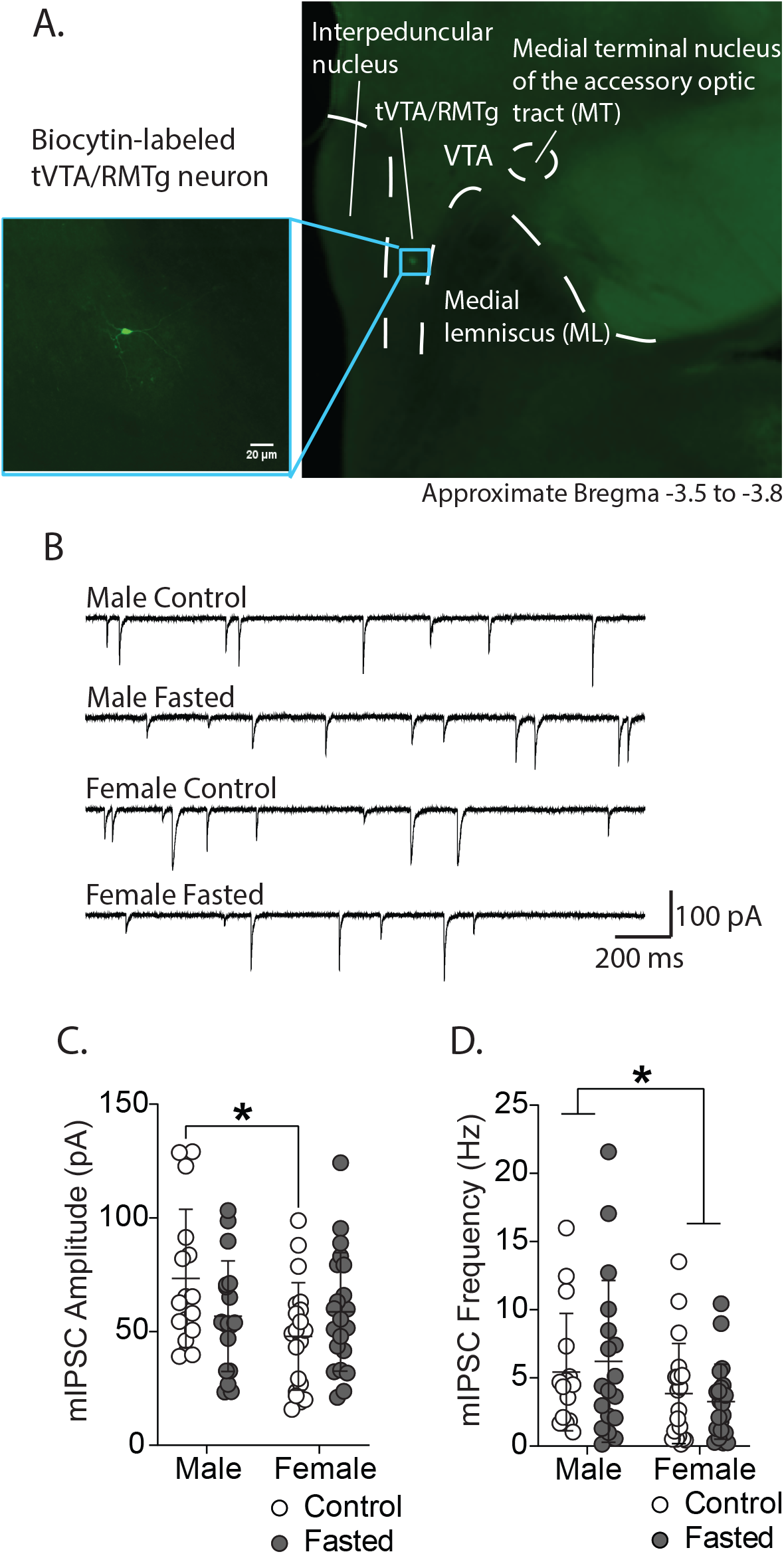
Effects of fasting and sex on in miniature inhibitory postsynaptic currents onto tVTA/RMTg GABA neurons of male and female mice. **A,** Example of a biocytin labeled tVTA/RMTg neuron located in a horizontal hemisected slice containing the VTA and caudal-medially located tVTA/RMTg. **B**, representative examples of mIPSC recordings from male and female fasted and control mice. **C**, mIPSC amplitude of GABA neurons of male and female fasted and control mice. mIPSC amplitude is less in female controls than male controls. **D**, mIPSC frequency onto GABA neurons of male and female fasted and control mice. mIPSC frequency onto tVTA/RMTg neurons of female fasted and control mice was less than in male control and fasted mice.

## Results

To identify GABAergic neurons in the tVTA/RMTg for electrophysiological recordings, we crossed VGAT^cre^ mice with Rosa-td Tomato mice. The VTA contains a heterogenous population of neurons, including neurons that contain and release dopamine, GABA, and glutamate (Nair-Roberts *et al*., 2008; Chieng *et al*., 2011). Caudally to the VTA, the neuronal population is primarily GABAergic (Barrot *et al*., 2012; Bourdy & Barrot, 2012). To validate our VGAT^cre^TdTomato mouse line, we examined colocalization between TdTomato fluorescence, a marker for GABAergic neurons and tyrosine hydroxylase (TH), a marker for dopamine neurons. Of all neurons expressing Td-Tomato, 2.6 ± 6.7 % co-express TH, with the remaining 97.3 ± 6.7 % expressing only Td-Tomato (Figure 1A,B), suggesting that the fluorescent label is expressed in non-dopaminergic neurons primarily medial to the medial lemiscus in the VTA. We focused our recordings to the VGAT^cre^TdTomato expressing neurons in caudal VTA slices, consistent with the GABAergic population of the tVTA/RMTg (Barrot *et al*., 2012).

### Effects of fasting on bodyweight and water consumption

To determine the effects of a 16 h overnight fast on VGAT^cre^Td-Tomato mice, we measured changes in body weight as well as food and water consumption (Figure 1C). Baseline body weights measured prior to the 16 h overnight fast were not different between fasted and control groups (fasting effect: F (1, 75) = 0.01, P = 0.9), but different between sexes (sex effect: F (1, 75) = 117.1, P<0.0001; male control (n = 18): 31.8 ± 3.5 *g*, male fasted (n = 20): 31.4 ± 4.1 g, female control (n = 21): 23.0 ± 2.5 g, female fasted (n = 20): 23.2 ± 3.7 g; Figure 1D). Similar to a previous study (Godfrey & Borgland, 2020), female control mice consumed more energy than males during the 16h period (t(37)=5.5, P<0.0001; male control (n = 18): 0.5 ± 0.09 kcal g^−1^; female control (n = 21): 0.7 ± 0.1 kcal g^−1^; Figure 1E). Fasting induced significant weight loss (fasting effect: F (1, 75) = 703.7, P<0.0001; male control (n = 18): 0.6 ± 2.03 %; female control (n = 21): 2.3 ± 2.4 %; male fasted (n = 20): −10.0 ± 1.6 %; female fasted (n = 20): −11.4 ± 2.0 %; Figure 1F). Weight loss was also affected by sex (sex effect: F (1, 75) = 0.1, P=0.7) with both male (P<0.0001) and female (P<0.0001) fasted mice losing more weight than their controls, as indicated by Sidak’s multiple comparisons test (sex x fasting interaction: (F(1, 75) = 11.0, P=0.0014). There was a main effect of fasting on water consumption, with fasted mice consuming less water than controls (fasting effect: F (1, 75) = 408.3, P<0.0001; male control (n = 18): 3.0 ± 0.6 mL; female control (n = 21): 3.3 ± 0.4 mL; male fasted (n = 20): 1.0 ± 0.4 mL; female fasted (n = 20): 1.1 ± 0.5 mL; Figure 1G). However, there was no sex difference in water consumption (sex effect: F (1, 75) = 2.2, P=0.1). Taken together, a 16h fast induces weight loss in male and female mice.

### Effects of fasting on tVTA/RMTg GABA neurons

To test the effect of fasting on GABAergic synaptic transmission onto tVTA/RMTg GABAergic neurons, we recorded miniature inhibitory postsynaptic currents (mIPSCs). There were no main effects of fasting or sex (fasting effect: F (1, 71) = 0.2, P=0.6; sex effect: F (1, 71) = 3.8, P=0.05). However, we observed a significant cross-over interaction between fasting and sex on mIPSC amplitude (sex x fasting interaction: F (1, 71) = 5.15, P=0.03; male control (N/n = 16/4): 73.4 ± 30.4 pA, female control (N/n = 20/4): 47.9 ± 23.6 pA; male fasted (N/n = 18/5): 56.8 ± 24.3 pA; female fasted (N/n = 21/4): 58.7 ± 26.0 pA; Figure 2C). A Sidak’s posthoc test indicates there was a significant difference between male and female control mIPSC amplitude (P= 0.02). To explore the GABAergic release onto tVTA/RMTg GABAergic neurons, we next examined mIPSC frequency. A main effect of sex was observed on mIPSC frequency (sex effect: F (1, 71) = 5.3, P=0.02; male control (N/n = 16/4): 5.4 ± 4.3 Hz; female control (N/n = 20/4): 3.8 ± 3.7 Hz; male fasted (N/n = 18/5): 6.2 ± 5.9 Hz; female fasted (N/n = 21/4): 3.3 ± 2.7 Hz; Figure 2D). However, there was no effect of fasting (fasting effect: F (1, 71) = 0.009, P=0.9) or sex x fasting interaction (Interaction: F (1, 71) = 0.5, P=0.5). Taken together, these results suggest that there were sex differences in GABAergic synaptic transmission onto tVTA/RMTg GABAergic neurons, but this is not strongly affected by fasting.

We next examined how fasting alters the excitability of tVTA/RMTg GABAergic neurons using current steps from −75 pA to 200 pA in 25 pA increments at 0.1 Hz while neurons were initially held at −70 mV. Analysis of the frequency-current (F-I) plot revealed a main effect of current steps (current effect: F (7.0, 378.0) = 21.3; P<0.0001) and a main effect of group (group effect: F (3.0, 378.0) = 5.6, P=0.0009, Figure 3C) with both male (P=0.02) and female (P=0.03) fasted being less excitable than male and female controls, as indicated by a Tukey’s multiple comparisons test. There was no effect of fasting or sex on the input resistance (fasting effect: F (1, 35) = 0.1, P=0.7; sex effect: F (1, 35) = 0.7, P=0.4; male control (N/n = 10/4): 0.6 ± 0.3 mOhms; female control (N/n = 9/3): 0.6 ± 0.3 mOhms, male fasted (N/n = 12/5): 0.7 ± 0.2 mOhms; female fasted (N/n = 8/4): 0.6 ± 0.4 mOhms; Figure 3D). However, a main effect of fasting was observed in the resting membrane potential (fasting effect: F (1, 42) = 4.8, P=0.03; male control (N/n = 9/6): −69.7 ± 3.4 mV; female control (N/n = 12/5): −69.7 ± 7.3 mV; male fasted (N/n = 11/6): −66.3 ± 7.6 mV; female fasted (N/n = 14/8): −64.9 ± 5.8 mV; Figure 3E) with fasted mice having a more depolarized resting membrane potential than controls. Taken together, these data suggest that fasting induces changes in excitability as well as resting membrane potential in both sexes.

**Figure 3.**
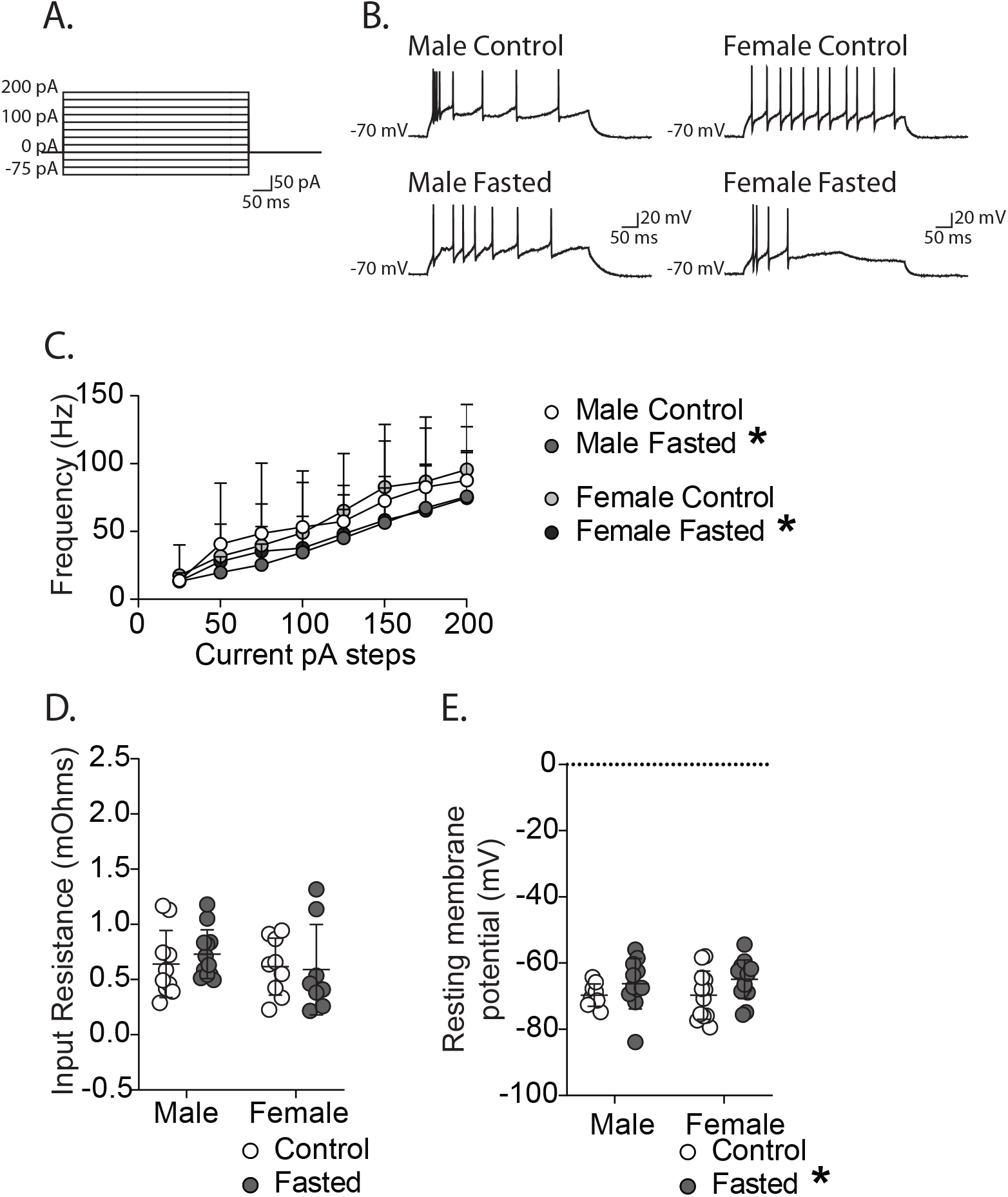
Effect of sex and fasting on excitability and intrinsic membrane properties of tVTA/RMTg GABA neurons. **A**, current step waveform of the 500 ms current steps. **B**, representative example recordings of tVTA/RMTg GABA neurons firing during a 100 pA step. **C**, frequency-current plot demonstrating a decrease in excitability of tVTA/RMTg GABA neurons following fasting in male and female mice. **D**, input resistance (mOhms) of tVTA/RMTg neurons of male and female fasted and control mice. No change in males or females following fasting. **E**, resting membrane potential (mV) of male and female fasted and control mice. tVTA/RMTg GABA neurons are more depolarized following fasting in both males and females.

### Effect of fasting on inhibitory synaptic transmission at LH GABA to tVTA/RMTg GABA synapses

We noted a significant amount of variability in the mIPSC recordings from control and fasted mice and therefore questioned if possible fasting-induced changes in GABAergic synaptic transmission may be driven by a specific input. LH GABAergic neurons become active during appetitive and consummatory behaviours (Jennings et al., 2015) and activation of LH GABA inputs to caudomedial VTA GABAergic neurons increases motivated behaviour for food or social reward (Nieh et al. 2015, 2016). Therefore, we predicted LH GABAergic inputs to the tVTA/RMTg may be susceptible to changes induced by fasting. We first verified the expression of ChR2 in LH GABAergic neurons (Figure 4A). We then recorded action potentials of LH GABAergic neurons evoked by optical stimulation (470 nm). In current clamp, action potentials were reliably evoked in LH GABAergic neurons with 5 Hz, 10 Hz, and 20 Hz optical stimulation (Figure 4B). Similarly, in voltage clamp of tVTA/RMTg GABAergic neurons, IPSCs were reliably evoked by 5 Hz, 10 Hz, and 20 Hz optical stimulation (Figure 4C). A 1 second stimulation produced a sustained depolarization in current clamp of LH GABAergic neurons, whereas this optical stimulation produced a single IPSC at tVTA/RMTg GABAergic neurons (Figure 4D, E). We next tested the nature of the LH GABA to tVTA/RMTg GABA input (Figure 4F). Optical stimulation of LH inputs evoked IPSCs onto tVTA/RMTg GABAergic neurons and this effect was blocked by picrotoxin (t(4)=4.4; baseline: −376.7 ± 191.1 pA, picrotoxin: −8.1 ± 5.3 pA, N/n = 5/2; Figure 4G). To test if this projection was monosynaptic, we applied the sodium channel blocker, TTX and then assessed if the current returned in the presence of 4-AP. TTX blocked optically evoked IPSCs (oIPSCs; baseline: −781.1 ± 317.9 pA, TTX: −39.6 ± 48.2 pA, N/n = 5/3), and this was restored by 4-AP (−741.4 ± 493.2 pA, N/n = 5/3; one-way ANOVA: (F (1.7, 6.7) = 11.3, P=0.008; Dunnett’s multiple comparisons tests baseline vs. TTX: P=0.009; baseline vs. TTX + 4-AP: P = 0.9; Figure 4H). Taken together, these data suggest that LH GABAergic neurons functionally synapse onto tVTA/RMTg GABAergic neurons though a monosynaptic projection.

**Figure 4.**
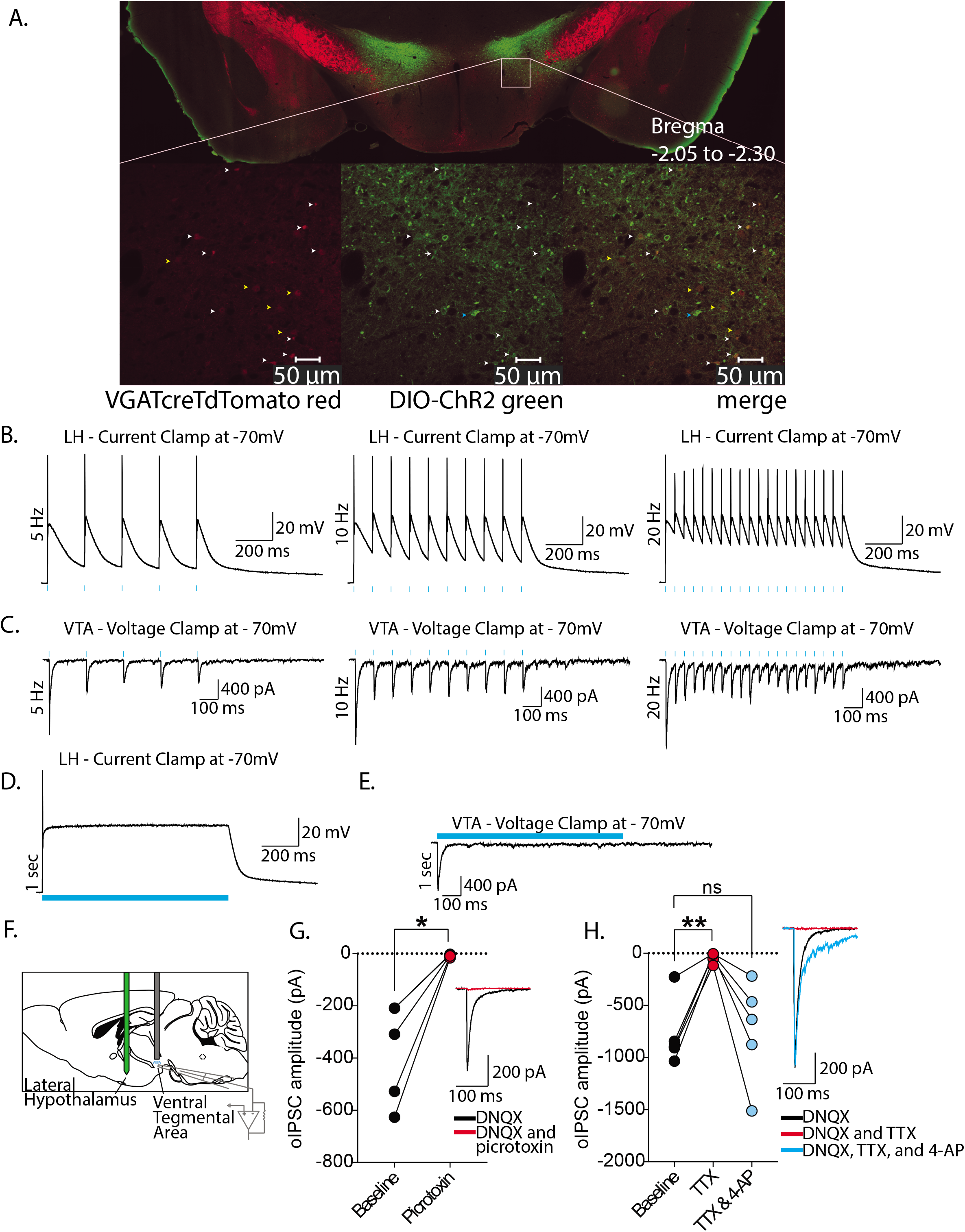
Validation of ChR2 in LH GABA neurons and their projection to tVTA/RMTg GABA neurons. **A,** slide scanner and confocal images of cells expressing VGAT^cre^TdTomato (red), DIO-ChR2 (green), and merged. Arrows refer to VGAT^cre^TdTomato neurons expressing ChR2 (white), VGAT^cre^TdTomato neurons not expressing ChR2 (yellow), or ChR2 neurons that are not VGAT^cre^TdTomato (blue). **B**, representative example recordings of optically stimulated action potentials in LH GABA neurons at 5 Hz, 10 Hz, and 20 Hz. **C**, representative example recordings of optically stimulated currents in tVTA/RMTg GABA neurons at 5 Hz, 10 Hz, and 20 Hz. **D**, representative example recording of the change in membrane potential during a 1 s optical stimulation of a LH GABA neuron. **E**, representative example recording of an optically stimulated 1 s current in a tVTA/RMTg GABA neuron. **F**, schematic of a sagittal brain slice, demonstrating the location of the LH injection and recording of tVTA/RMTg GABA neurons. **G,** optically evoked currents in tVTA/RMTg GABA neurons are blocked with bath application of picrotoxin. **H**, optically evoked currents of tVTA/RMTg GABA neurons are blocked with bath application of TTX, and then restored following bath application of TTX and 4-AP.

To probe the quantal properties of LH GABAergic input to tVTA/RMTg GABA neurons, extracellular Ca^2+^ was replaced by Sr^2+^ leading to asynchronous exocytosis of vesicles allowing the resolution of quantal synaptic events. Under these experimental conditions, optogenetic stimulation of LH GABAergic inputs to tVTA/RMTg GABA neurons evoked asynchronous (as) release events (asIPSCs). In the first 50 ms, the optical stimulation was frequently followed by a summation of synchronized events that interfere with identification of asIPSCs. Hence, we followed previously established protocols and analyzed 50 to 350 ms following the optical stimulation for asIPSCs (MacAskill *et al*., 2014; Geddes *et al*., 2016). We also identified asIPSCs between 700 to 1000 ms following the optical stimulation, a period where the effects of the optical stimulation have mostly ended. This gave us time periods to examine both the LH GABA input (50 to 350 ms) and all inputs (700 to 1000 ms). Between 50 to 350 ms following the optical stimulation, there was a main effect of fasting (fasting effect: F (1, 47) = 7.1, P=0.01) and a main effect of sex (sex effect: F (1, 47) = 5.5, P=0.02) on asIPSC amplitudes (male control (N/n = 14/4): 76.4 ± 34.2 pA; female control (N/n = 13/5): 57.4 ± 22.9 pA, male fasted (N/n = 13/4): 55.3 ± 13.5 pA; female fasted (N/n = 11/5): 43.2 ± 16.1 pA; Figure 5B). However, between 700 to 1000 ms following the optical stimulation there was neither a main effect of fasting (fasting effect: F (1, 47) = 0.02, P=0.9) or sex (sex effect: F (1, 47) = 0.001, P=1.0) on asIPSC amplitude (male control (N/n = 14/4): 48.2 ± 18.3 pA; female control (N/n = 13.5): 49.4 ± 17.3 pA, male fasted (N/n = 13/4): 50.4 ± 26.7 pA; female fasted (N/n = 11/5): 48.9 ± 11.6 pA; Figure 5C).

**Figure 5.**
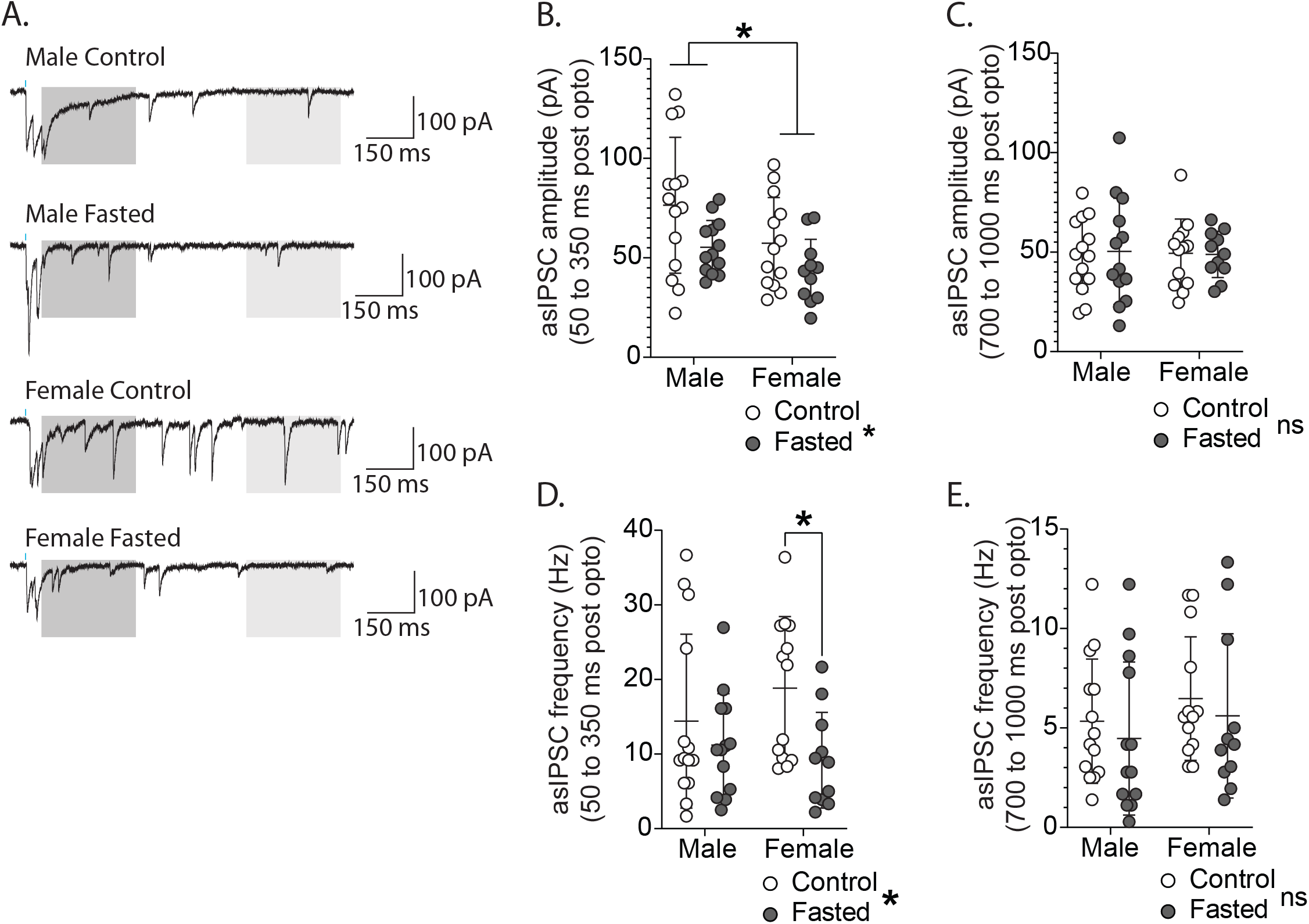
Strontium-induced asynchronous IPSCs at LH GABA to tVTA/RMTg GABA synapses. **A,** representative example recordings of optically induced asynchronous asISPCs at LH GABA to VTA GABA synapse of male and female fasted and control mice. **B**, optically evoked asIPSC amplitude (pA) at LH GABA to tVTA/RMTg GABA synapses from 50 to 350 ms following optical stimulation. The amplitude of asIPSCs decreased following fasting in both male and female mice. The asIPSC amplitude was also smaller in females compared to males in both fasted and control mice. **C,** optically evoked asIPSC amplitude (pA) at LH GABA to tVTA/RMTg GABA synapses from 700 to 1000 ms following optical stimulation. Fasting does not affect the asIPSC amplitude during this time period. **D**, asIPSC frequency (Hz) at LH GABA to tVTA/RMTg GABA synapses from 50 to 350 ms following optical stimulation. The frequency of asIPSCs decreased following fasting in female mice. **E**, asIPSC frequency (Hz) at LH GABA to tVTA/RMTg GABA synapses from 700 to 1000 ms following optical stimulation. Fasting does not affect the asIPSC amplitude during this time period.

Frequency of asIPSCs between 50 to 350 ms following the optical stimulation was significantly different in fasted mice (Fasting effect: F(1, 47) = 6.4, P=0.02) but no effect of sex (sex effect: F (1, 47) = 0.2, P=0.6), or a sex x fasting interaction (F (1, 47) = 1.6, P=0.2). Given that we observed sex differences in mIPSC frequency and that asIPSC frequency appeared to be reduced in both male and female mice preventing a sex x fasting interaction, we made the a priori hypothesis that fasting would affect asIPSC frequency differently in males and females. A Sidak’s multiple comparisons test on the effect of fasting revealed no difference in frequency between male control (14.4 ± 11.7 Hz, N/n = 14/4) and fasted mice (11.2 ± 6.9 Hz, N/n = 13/4; P=0.6), but a significant difference in female control (18.9 ± 9.6 Hz, N/n = 13/5) and fasted mice (9.2 ± 6.4 Hz, N/n = 11/5; P=0.02; Figure 5D). Between 700 to 1000 ms following optical stimulation, there was no main effect of fasting (F (1, 47) = 0.8, P=0.4), sex effect (F (1, 47) = 1.3, P=0.3), or sex x fasting interaction on asIPSC frequency (interaction: F (1, 47) = 0.00003, P=1.0; male control (N/n = 14/4): 5.4 ± 3.1 Hz; female control (N/n = 13/5): 6.5 ± 3.1 Hz; male fasted (N/n = 13/4): 4.5 ± 3.8 Hz; female fasted (N/n = 11/5): 5.6 ± 4.1 Hz; Figure 5E). Taken together, these data indicate that fasting decreases amplitude of asIPSCs induced by optical stimulation of LH GABA inputs to tVTA/RMTg GABA in all mice, but only decreases asIPSC frequency in female mice.

To characterize properties of short-term synaptic plasticity at these synapses, we next examined if oIPSCs from LH GABAergic inputs to tVTA/RMTg GABAergic neurons were altered by fasting using a 20Hz optical train stimulation (Figure 6A-C). We first assessed for changes in vesicle release kinetics determined by the readily releasable pool size (RRP_train_) and steady-state oIPSC amplitude, reflecting the replenishment of the RRP after it has been depleted (Thanawala & Regehr, 2013, 2016). Analysis of the first oIPSC revealed an effect of fasting on the amplitude of oIPSCs onto tVTA/RMTg GABAergic neurons (fasting effect: F (1, 79) = 13.1, P=0.0005; male control (N/n = 21/5): −553.2 ± 462.0 pA; male fasted (N/n = 23/5): −281.4 ± 230.0 pA; female control (N/n = 19/5): −640.3 ± 461.7 pA; female fasted (N/n = 20/5): −331.1 ± 251.3 pA; Figure 6D), but no effect of sex (sex effect: F (1, 79) = 0.7, P=0.4). There was no significant interaction of sex x fasting (F (1, 79) = 0.05, P=0.8). In addition, there was a fasting effect on the RRP_train_ (fasting effect: F (1, 75) = 10.3, P=0.002; male control (N/n = 20/5): 1186 ± 1121 pA, male fasted (N/n = 23/5): 685.8 ± 615.5 pA, female control (N/n = 18/5): 1692 ± 1356 pA, female fasted (N/n = 18/5): 783.1 ± 656.6 pA; Figure 6E,F) and the steady-state oIPSC amplitude (fasting effect: F (1, 79) = 7.3, P=0.009 pA; male control (N/n = 21/5): 241.4 ± 207.2 pA, male fasted (N/n = 23/5): 99.2 ± 82.0 pA, female control (N/n = 19/5): 205.8 ± 193.9 pA, female fasted (N/n = 20/5): 157.6 ± 136.0 pA; Figure 6G). There was no effect of sex or sex x fasting interaction in either the RRP_train_ (sex effect: F (1, 75) = 1.9, P=0.2; fasting x sex: F (1, 75) = 0.9, P=0.4) or the steady-state oIPSC amplitude (sex effect: F (1, 79) = 0.1, P=0.7; fasting x sex: F (1, 79) = 1.8, P=0.2). Thus, fasting decreased the amplitude of the first pulse, the effective RRP size, and the steady-state oIPSC amplitude of the LH GABAergic input to tVTA/RMTg GABAergic neurons in male and female mice.

**Figure 6.**
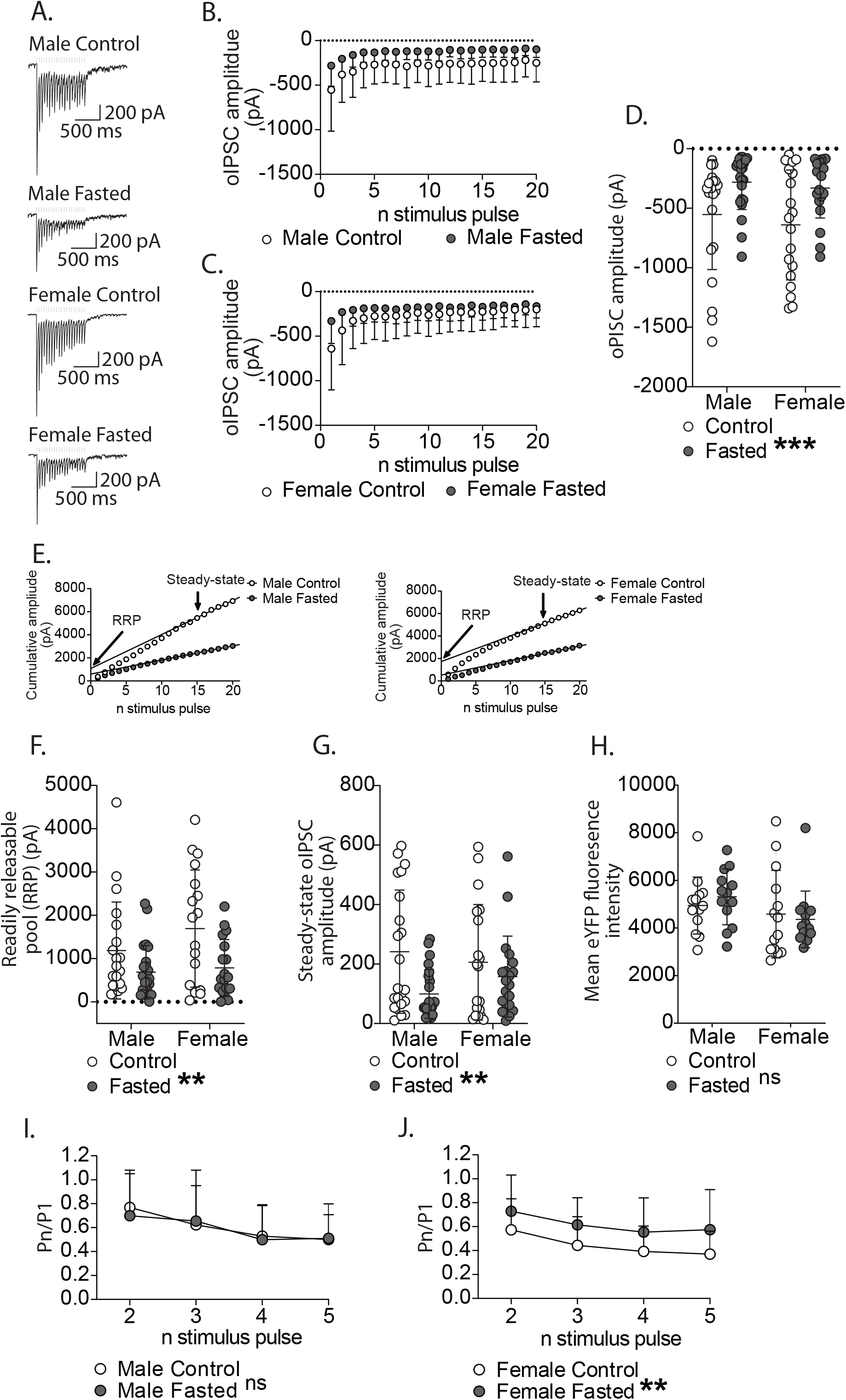
Short-term synaptic plasticity at tVTA/RMTg GABA synapses after an optically induced 20 Hz train stimulation of LH GABAergic terminals. **A,** representative example recordings of optically induced currents at 20 Hz at LH GABA to tVTA/RMTg GABA synapses of male and female fasted and control mice. **B**, time course showing the average oIPSC amplitude for all 20 pulses in male fasted and control mice. **C,** time course showing the average oIPSC amplitude for all 20 pulses in female fasted and control mice. **D,** oIPSC amplitude (pA) of the first pulse in the 20 Hz stimulation. oIPSC amplitude decreases following fasting in both male and female mice. **E**, representative example of the cumulative amplitude, used to calculate the readily releasable pool (RRP_train_, pA) and the steady state oIPSC amplitude (pA), from recordings of a male and female fasted and control tVTA/RMTg GABA neurons. **F**, the RRP_train_ and steady state oIPSCs (pA) at LH GABA to tVTA/RMTg GABA synapses in male and female control and fasted mice. The RRP_train_ decreases following fasting in both male and female mice. **G,** the steady-state oIPSC amplitude at LH GABA to tVTA/RMTg GABA synapses in male and female control and fasted mice. The steady-state oIPSC amplitude decreases following fasting in both male and female mice. **H**, the mean eYFP fluorescence intensity in male and female control and fasted mice. The eYFP fluorescence intensity is not different between control or fasted male or female mice. **I**, Pn/P1 plotted for the first 5 pulses of the optically induced 20 Hz train stimulation in male control and fasted mice. A paired pulse depression was observed in both male control and fasted mice. **J**, Pn/P1 plotted for the first 5 pulses of the optically induced 20 Hz train stimulation in female control and fasted mice. A paired pulse depression was observed in both female control and fasted, but there was less of a depression following fasting.

To determine if there are fasting-induced differences in short-term plasticity, we measured the ratio of the amplitude of oIPSCs (Pn) over the first oIPSC (P1) for each of the first 5 pulses. While male mice exhibited a paired-pulse depression (pulse effect: F(2.3,94.9) = 23.4, P<0.0001), there was no effect of fasting (fasting effect: F(1,42) = 0.03, P=0.9; Figure 6I). However, there was a significant paired pulse depression in female mice (pulse effect: F(2.8, 104.9) = 18.6, P<0.0001), and an effect of fasting (fasting effect: F(1,37) = 5.09, P=0.03; Figure 6J). Taken together, fasting alters the release probability of LH GABA inputs of female mice.

To address the potential of differences in ChR2 expression between groups, we quantified eYFP fluorescence intensity from slices used for the 20 Hz train optical stimulation recordings. There were no fasting effects (fasting effect: F (1, 51) = 0.04, P=0.8) or sex effects on eYFP fluorescence intensity (sex effect: F (1, 51) = 3.07, P=0.08; fasting x sex: F (1, 51) = 0.6, P=0.4); male control (N/n = 13/5): 4943 ± 1197, male fasted (N/n = 13/5): 5314 ± 1174, female control (N/n = 14/5): 4583 ± 1839, female fasted (N/n = 15/5): 4365 ± 1190; Figure 6H). Furthermore, we did not observe any differences between groups in the number of neurons recorded that received ChR2 input (Male control: 27/37, 73%; Male fasted: 29/40, 73%, Female control: 25/37, 68%; Female fasted: 30/43, 70%). Thus, presumably ChR2 expression at LH GABA inputs to the VTA is similar between groups and not driving electrophysiological differences.

### Effect of fasting on firing rate of tVTA/RMTg GABA neurons during optical stimulation of LH GABA inputs

We next tested whether the effects of a 20 Hz optical stimulation on synaptic transmission of LH GABAergic terminals influenced evoked firing of tVTA/RMTg GABAergic neurons. We analyzed firing properties of tVTA/RMTg GABAergic neurons over the entire 1 s step, as well as just the first 200 ms, which is the duration of 5 pulses at 20 Hz. In males, over the 1s step, there was no effect of optical stimulation or fasting on number of action potentials (stimulation effect: F (1, 22) = 0.2, P=0.9; fasting (fasting effect: F (1, 22) = 0.002, P=0.9; fasting x stimulation: F (1, 22) = 0.2, P=0.7; male control pre-OPTO (N/n = 12/6): 14.0 ± 16.1, male fasted pre-OPTO (N/n = 12/6): 12.9 ± 9.3, male control OPTO (N/n = 12/6): 12.3 ± 13.2, male fasted OPTO (N/n = 12/6): 12.9 ± 12.8; Figure 7A-D). Within the first 200 ms of the current step, there was an effect of optical stimulation on action potentials in males (stimulation effect: F (1, 22) = 17.5, P=0.0004; fasting effect: F (1, 22) = 0.3, P=0.6; fasting x stimulation interaction: F (1, 22) = 1.4, P=0.2; male control pre-OPTO (N/n = 12/6): 6.7 ± 4.7, male fasted pre-OPTO (N/n = 12/6): 5.3 ± 4.6, male control with OPTO (N/n = 12/6): 4.8 ± 4.3, male fasted with OPTO (N/n = 12/6): 4.3 ± 4.3; Figure 7E). In female mice, over the 1s step there was an effect of optical stimulation on action potentials (stimulation effect: F (1, 20) = 9.5, P=0.006; female control pre-OPTO (N/n = 10/5): 26.0 ± 35.2, female fasted pre-OPTO (N/n = 12/7): 9.2 ± 7.0, female control with OPTO (N/n = 10/5): 20.0 ± 34.5, female fasted with OPTO (N/n = 12/7): 8.5 ± 8.6; Figure 7F-H). However, like male mice, there was no effect of fasting on action potentials in female mice (fasting effect: F (1, 20) = 1.9, P=0.2; fasting x stimulation: F (1, 20) = 6.0, P=0.02). There were no significant differences between control and fasted in pre-OPTO (P = 0.2) or OPTO (P = 0.5) groups (Figure 7F-H). Within the first 200 ms, there was an effect of stimulation in females (stimulation effect: F (1, 20) = 17.7, P=0.004; female control pre-OPTO (N/n = 10/5): 9.3 ± 7.2, female fasted pre-OPTO (N/n = 12/7): 3.9 ± 3.1, female control OPTO (N/n = 10/5): 5.7 ± 7.0, female fasted OPTO (N/n = 12/7): 3.3 ± 3.2; Figure 7I). There was a significant interaction (fasting x stimulation: F (1, 20) = 8.2, P=0.009), but no effect of fasting (fasting effect: F (1, 20) = 3.04, P=0.09). A Sidak’s posthoc test indicated a significant effect of fasting during pre-OPTO (p = 0.04), but not OPTO (p = 0.5). Taken together, optical stimulation of LH GABAergic input decreases firing of tVTA/RMTg GABAergic neurons and fasting decreases the number of action potentials in the first 200 ms but this effect is lost after optogenetic stimulation.

**Figure 7.**
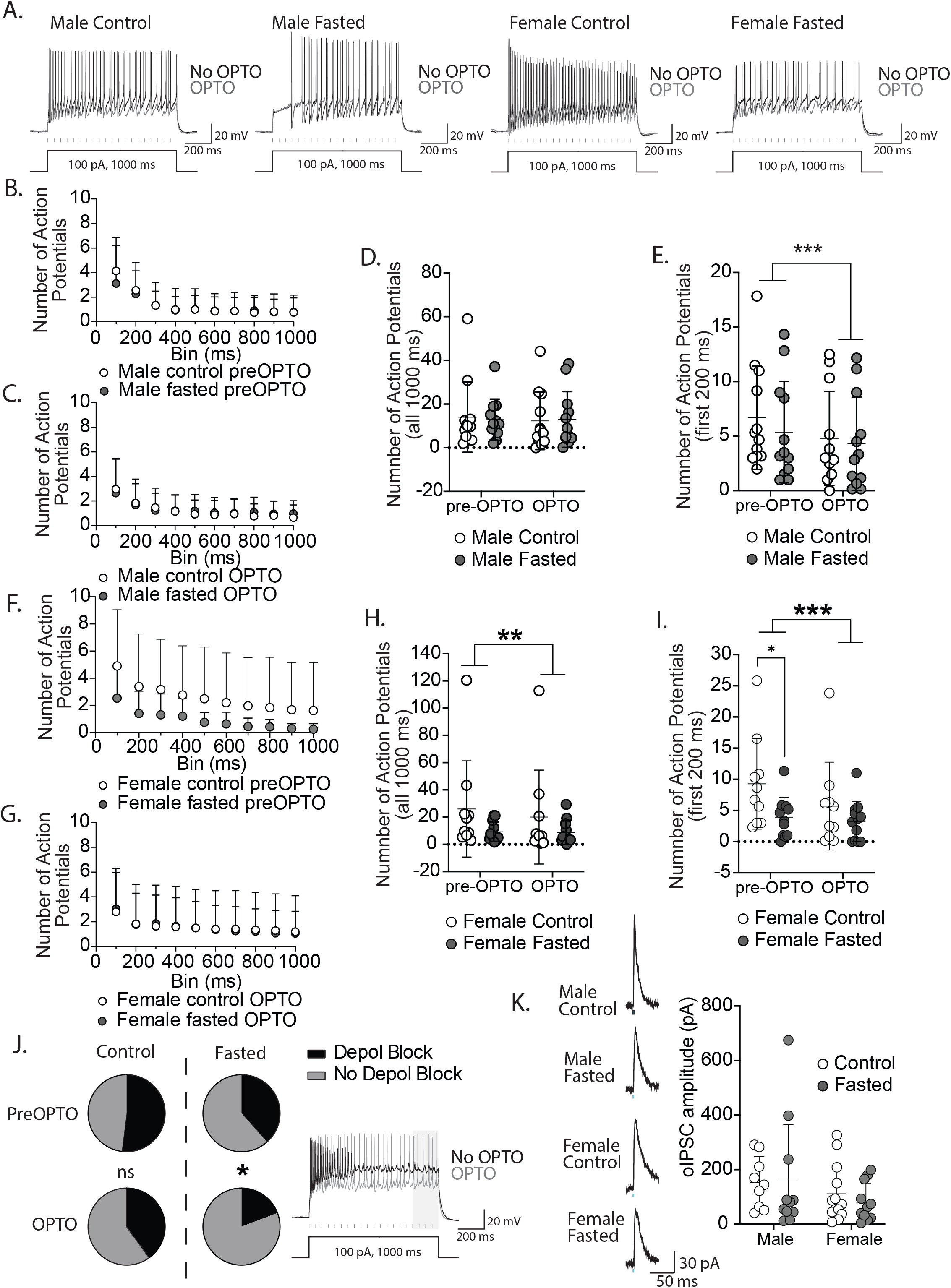
Effects of optical stimulation of the LH GABA input on the firing of tVTA/RMTg GABA neurons. **A,** representative example recordings of action potentials from tVTA/RMTg GABA neurons induced through a 1 s, 100 pA current step, before (black) and with (grey) an optically induced 20 Hz stimulation of LH GABA terminals in male and female fasted and control mice. **B**, a time course over the 1 s current step of the average number of action potentials in each 100 ms bin in male control and fasted before the optical stimulation. **C**, a time course over the 1 s current step of the average number of action potentials in each 100 ms bin in male control and fasted with the optical stimulation. **D**, average number of action potentials during the 1 s current step in male mice before and with optical stimulation of LH GABA terminals in the tVTA/RMTg. Optical stimulation of the LH GABA neuron terminals in the tVTA/RMTg did not affect the average number of action potentials during the complete 1 s current step. **E**, average number of action potentials during the first 200 ms of the current step in male mice before and during optical stimulation. The average number of action potentials during the first 200 ms of the current step decreased with optical stimulation of LH GABA neuron terminals in the tVTA/RMTg. **F**, a time course over the 1 s current step of the average number of action potentials in each 100 ms bin in female control and fasted before the optical stimulation. **G**, a time course over the 1 s current step of the average number of action potentials in each 100 ms bin in female control and fasted during optical stimulation. **H**, average number of action potentials during the 1 s current step in female mice before and during optical stimulation. The average number of action potentials in the 1 s current step decreased with the optical stimulation. **I**, average number of action potentials during the first 200 ms of the current step in female mice before and during optical stimulation. The average number of action potentials in the first 200 ms of the current step decreases with optical stimulation. **J**, The proportion of cells with depolarization block during the 1s 100 pA current step in control and fasted, and pre-OPTO and OPTO. Following fasting, the number of cells with depolarization block decreased when the 1 s current step was paired with the optical stimulation. **K**, Optically evoked outward GABA_A_ currents recorded at −40 mV in K^+^Gluconate (low Cl^-^) internal solution from control (open circles) or fasted (filled circles) mice. Inset, example traces from male and female control and fasted mice.

We noticed that depolarization block occurred in a significant proportion of neurons with a 1 s current step, defined as no action potentials in the final 200 ms of the current step. Prior to optical stimulation, we found no difference in the occurrence of depolarization block between male and female control mice (male control (N/n = 12/6): 50 %, female control (female control (N/n = 13/5): 53.8 %; P >0.9999). Therefore, we grouped male and female mice together to examine the effects of fasting on optogenetic stimulation on the proportion of neurons with depolarization block. In control mice, there was no difference between pre stimulation (control pre-OPTO (N/n = 25/11): 52%) and optical stimulation (control OPTO (N/n = 25/11): 40%; P = 0.2380; Figure 7J). However, in fasted mice, optical stimulation significantly decreased the proportion of neurons with depolarization block (fasted pre-OPTO (N/n = 26/13): 38.5 %, fasted OPTO (N/n = 26/13): 19.2%; P = 0.04; Figure 7J). These data suggest that during fasting, the function of the LH GABA to tVTA/RMTg GABA synapse is preserved under depolarized conditions.

We recorded our voltage clamp experiments evoking IPSCs in the presence of high Cl^-^ in our internal solution whereas current clamp experiments to evoke firing were performed in the presence of K^+^ gluconate. Therefore, to confirm that we were indeed inducing GABAergic currents sufficient to alter tVTA/RMTg neuronal excitability, we plotted amplitudes of GABA currents evoked at −40 mV in voltage clamp. Average oIPSC amplitudes were 156 ± 4 pA (n = 19) for males and 96 ± 21 pA (n = 22) for females (Figure 7K). There was no sex x fasting interaction (F(1,40) = 0.21, P = 0.65) or main effects of sex (F(1,40) = 2.3, P = 0.13) or fasting (F(1,40) = 0.1, P = 0.7). Thus, during depolarizing current steps, it is likely that sufficient GABA_A_ currents could modulate firing activity.

We next examined how optical stimulation influenced action potential characteristics. Consistent with a reduction in action potential number, optical stimulation of LH GABA inputs increased the latency to fire in both males (stimulation effect: F (1, 21) = 34.3, P<0.0001; male control pre-OPTO (N/n = 11/6): 20.4 ± 24.2 ms, male fasted pre-OPTO (N/n = 12/6): 50.3 ± 62.4 ms, male control OPTO (N/n = 11/6): 35.9 ± 35.6 ms, male fasted OPTO (N/n = 12/6): 71.7 ± 76.6 ms; Figure 8A) and females (stimulation effect: F (1, 20) = 5.4, P=0.03; female control pre-OPTO (N/n = 10/5): 52.6 ± 79.4 ms, female fasted pre-OPTO (N/n = 12/7): 56.9 ± 82.4 ms, female control OPTO (N/n = 10/5): 125.5 ± 164.0 ms, female fasted OPTO (N/n = 12/7): 79.8 ± 109.6 ms; Figure 8B). No effect of fasting was observed in either males (fasting effect: F (1, 22) = 2.2, P=0.2, fasting x stimulation: F (1, 21) = 0.5, P=0.5) or females (fasting effect: F (1, 20) = 0.2, P=0.6; fasting x stimulation: F (1, 20) = 1.5, P=0.2). However, there was a significant effect of fasting on the percent change in latency to fire (fasting effect: F (1, 41) = 5.6, P=0.02), but no effect of sex or interaction (sex effect: F(1, 41) = 2.6, P=0.1, sex x fasting interaction: F(1, 41) = 0.002, P=0.9; Figure 8C). There was no change in AHP due to 20 Hz optical stimulation analyzed over the 1 s current step in either males (stimulation effect: F (1, 21) = 1.7, P=0.2; male control pre-OPTO (N/n = 11/6): 15.2 ± 3.7 mV, male control OPTO (N/n = 11/6): 15.2 ± 4.4 mV, male fasted pre-OPTO (N/n = 12/6): 17.3 ± 5.6 mV, male fasted OPTO (N/n = 12/6): 18.5 ± 5.7 mV; Figure 8D-F) or females (stimulation effect: F (1, 20) = 0.7, P=0.4; female control pre-OPTO (N/n = 10/5): 15.1 ± 5.6 mV, female control OPTO (N/n = 10/5): 16.2 ± 6.3 mV, female fasted pre-OPTO (N/n = 12/7): 16.7 ± 5.6 mV, female fasted OPTO (N/n = 12/7): 16.4 ± 5.4 mV; Figure 8H-J). There was no effect of fasting in the 1 s current step observed in either males (fasting effect: F (1, 21) = 1.7, P=0.2; fasting x stimulation: F (1, 21) = 1.4, P=0.2; Figure 8F) or females (fasting effect: F (1, 20) = 0.2, P=0.7; fasting x stimulation: F (1, 20) = 2.1, P=0.2; Figure 8J). When analyzing the first 200 ms of the current step, optical stimulation had no effect on the AHP in either males (stimulation effect: F (1, 21) = 2.1, P=0.2; male control pre-OPTO (N/n = 11/6): 14.6 ± 4.4 mV, male control OPTO (N/n = 11/6): 15.2 ± 4.4 mV, male fasted pre-OPTO (N/n = 12/6): 17.5 ± 6.0 mV, male fasted OPTO (N/n = 12/6): 18.5 ± 6.0 mV; Figure 8G) or females (stimulation effect: F (1, 18) = 0.5, P=0.5; female control pre-OPTO (N/n = 10/5): 15.7 ± 6.0 mV, female control OPTO (N/n = 10/5): 16.4 ± 6.5 mV, female fasted pre-OPTO (N/n = 12/7): 17.2 ± 5.8 mV, female fasted OPTO (N/n = 10/7): 16.1 ± 5.6 mV; Figure 8K). Furthermore, there was no effect of fasting on AHP observed in either males (fasting effect: F (1, 21) = 1.9, P=0.2; fasting x stimulation: F (1, 21) = 0.1, P=0.7) or females (fasting effect: F (1, 20) = 0.2, P=0. 6; fasting x stimulation: F (1, 18) = 0.7, P=0.4). Taken together, these data suggest that the decrease in firing of tVTA/RMTg GABA neurons due to optogenetic stimulation of LH GABA inputs is mediated by an increased latency to fire rather than a change in AHP.

**Figure 8.**
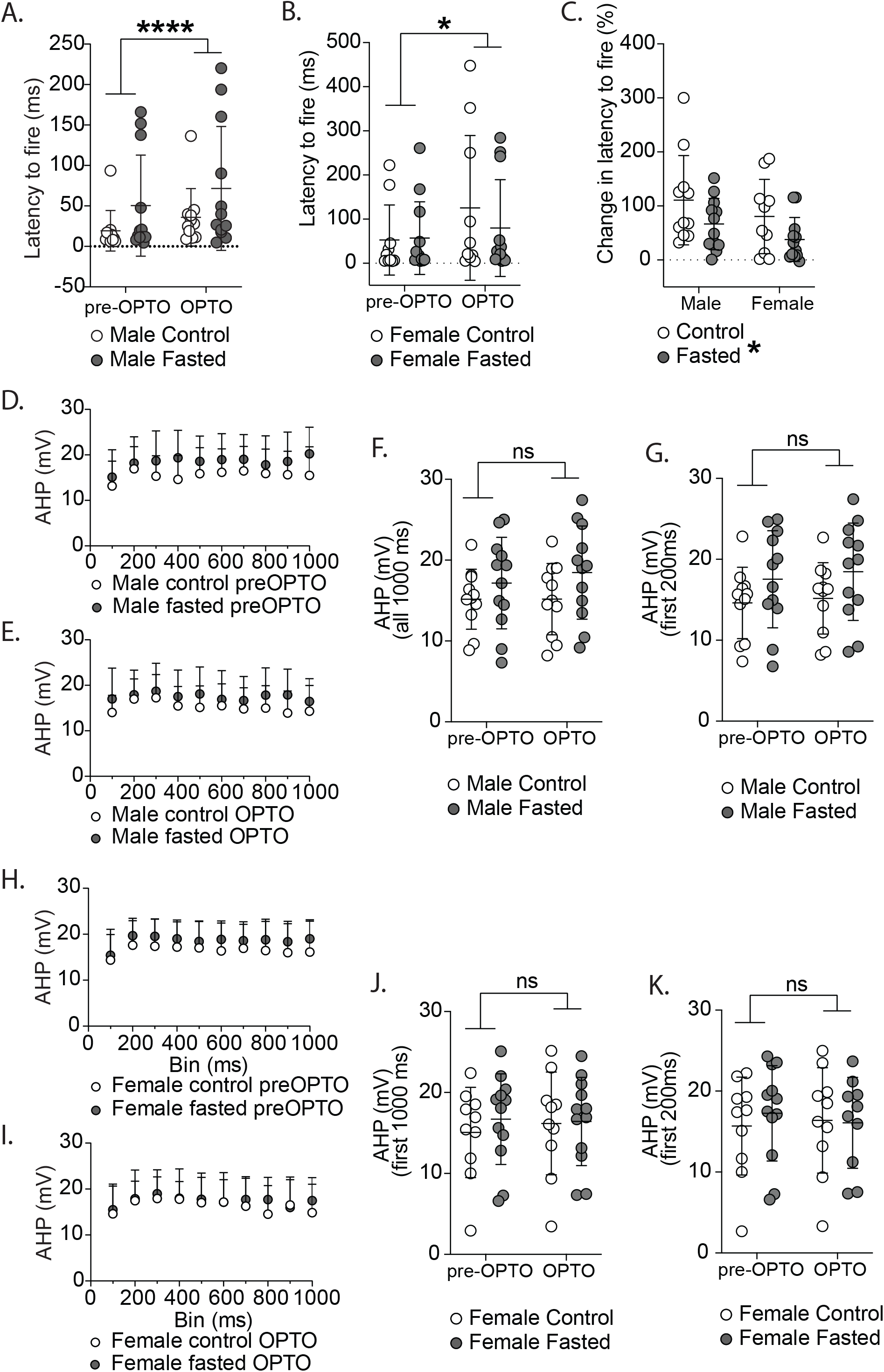
Effects of optical stimulation of the LH GABA input on action potential properties of tVTA/RMTg GABA neurons. **A**, latency to fire (ms) an action potential during the 1 s current step before and with optical stimulation of the LH input in male mice. The latency to fire was greater with optical stimulation in male mice. **B**, latency to fire (ms) an action potential during the 1 s current step before and with optical stimulation in female mice. The latency to fire was greater with the optical stimulation in female mice. **C**, The change in latency to fire (%) from LH GABA optical stimulation in male and female, fasted and control mice. Following fasting, the change in latency to fire (%) induced by optical stimulation of LH GABA terminals decreased. **D**, a time course over the 1 s current step of the average AHP (mV) in each 100 ms bin in male control and fasted before optical stimulation. **E**, a time course over the 1 s current step of the average AHP (mV) in each 100 ms bin in male control and fasted with the optical stimulation. **F**, the average AHP (mV) during the 1 s current step in male control and fasted, pre-OPTO and OPTO. The average AHP (mV) during the 1 s current step was not affected by optical stimulation in male mice. **G**, the average AHP (mV) during the first 200 ms of the current step in male control and fasted, pre-OPTO and OPTO. The average AHP during the first 200 ms of the current step was not affected by optical stimulation in male mice. **H**, a time course over the 1 s current step of the average AHP in each 100 ms bin in female control and fasted mice before optical stimulation. **I**, a time course over the 1 s current step of the average AHP in each 100 ms bin in female control and fasted mice with optical stimulation. **J**, the average AHP during 1 s current step in female control and fasted, pre-OPTO and OPTO. The average AHP during the 1 s current step was not affect by optical stimulation in female mice. **K**, the average AHP during the first 200 ms of the current step in female control and fasted, pre-OPTO and OPTO. The average AHP during the first 200 ms of the current step was not affected by optical stimulation in female mice.

## Discussion

Here, we found that fasting decreased the excitability of tVTA/RMTg GABAergic neurons, suggesting that this population of neurons is also sensitive to motivational state. LH GABAergic neurons make a monosynaptic projection to tVTA/RMTg GABAergic neurons. Inhibitory synaptic transmission of this input onto tVTA/RMTg GABAergic neurons was sensitive to fasting, such that fasting decreased the release probability at LH GABAergic terminals of female mice and decreased the amplitude of optically induced currents, RRP size, and RRP replenishment in both male and female mice. Finally, activation of the LH GABAergic input to the VTA decreased firing of tVTA/RMTg GABAergic neurons, and this effect was minimized in fasted mice. These results suggest that fasting can reduce the excitability of tVTA/RMTg GABAergic neurons and the inhibitory strength of the LH GABAergic input to these neurons. This may lead to the preservation of the function of this synapse. Because LH GABAergic neurons can disinhibit VTA dopamine neurons leading to increased NAc dopamine and behavioural activation (Nieh et al., 2016), activation of weaker LH GABA inputs, following fasting, may still be sufficient to induce suppression of tVTA/RMTg GABAergic neuronal activity, given the concurrent decrease in excitability of these neurons, to allow for foraging.

Consistent with our previous work (Godfrey & Borgland, 2020), a 16h overnight fast reduced body weight in male and female mice. Previous studies have reported that this reduction in weight is due to both a reduction in body weight (60%) and gastric emptying (40%) (Prior *et al*., 2012). Although we did not see sex differences in weight change, our previous findings demonstrated that, as a physiological stressor, fasting affects males and females differently, with females observed to have a higher concentration of corticosterone (Godfrey & Borgland, 2020). Furthermore, given that this duration of fast can increase food approach behaviours and food intake after fasting (Godfrey and Borgland, 2020), this likely represents a motivational state of hunger.

### Sex differences of GABAergic synaptic transmission onto tVTA/RMTg neurons

In unfasted control mice, we observed sex differences in inhibitory synaptic transmission onto tVTA/RMTg GABAergic neurons. Female mice had decreased amplitude and frequency of mIPSCs or asIPSCs onto tVTA/RMTg GABAergic neurons compared to male mice. This could be due to a decrease in postsynaptic GABA_A_ receptor current size leading to a decrease in mIPSC event detection or due to a decrease in number of inhibitory synapses onto tVTA/RMTg neurons in female mice. Notably, the delta-containing subunit of GABA_A_ receptors are regulated by ovarian hormones (Stell *et al*., 2003; Maguire *et al*., 2005). Associated *Gabrd* transcript levels in the VTA are altered across the estrous cycle (Melón *et al*., 2017) and δ-GABAA receptors can influence lateral diffusion of the receptor across the membrane and its ability to cluster at the synapse, ultimately influencing tonic inhibition (Jacob *et al*., 2008). Thus, it is possible this difference in inhibitory synaptic transmission could be due to either differential expression of δ-GABA_A_ receptors or their regulation by neurosteroids. Alternatively, sex differences in GABAergic synaptic transmission could be due to differences in endocannabinoid modulation of inhibitory inputs to tVTA/RMTg neurons. Notably, tonic endocannabinoid signaling at inhibitory inputs to dopamine neurons was greater in female compared to male rats (Melis *et al*., 2013) suggesting potential sex differences in endocannabinoid signaling within the VTA.

Although fasting did not alter inhibitory synaptic transmission of tVTA/RMTg neurons, fasting suppressed the excitability of tVTA/RMTg neurons of both male and female mice. These changes in excitability were not paralleled with changes in input resistance, suggesting that this change may be driven by changes in the strength of synaptic input rather than intrinsic conductances. It is possible that enhanced tonic GABA release may underlie the fasting-induced decrease in excitability of tVTA/RMTg neurons of male and female mice, as tonic inhibition strongly influences neuronal excitability (Farrant & Nusser, 2005). These effects were surprising as, unlike fasting, other aversive states such as pain, predator odours, opioid withdrawal and noxious stimuli increase activity of tVTA/RMTg neurons (Sánchez-Catalán *et al*., 2017; Li *et al*., 2019a). Notably, the resting membrane potential was more depolarized following fasting; a change that should theoretically produce an increase in excitability. It is possible that there are independent changes in excitatory input that could influence the resting membrane potential. For example, the lateral habenula (LHb) sends excitatory projections to tVTA/RMTg neurons leading to their activation during aversive stimuli (Brown *et al*., 2017; Li *et al*., 2019b).

### Sex differences of LH GABA inputs to tVTA/RMTg GABA synapses

LH GABAergic neurons functionally synapse onto tVTA/RMTg GABAergic neurons through a monosynaptic projection. This effect is consistent with previous studies that have recorded from VTA GABAergic interneurons that disinhibit dopamine neurons (Nieh et al., 2015). tVTA/RMTg GABAergic neurons send dense projections to dopamine neurons in the VTA and substantia nigra (SN) (Barrot *et al*., 2012; Bourdy & Barrot, 2012). Due to their dense projections and proximal dendritic connections, they have a strong influence over the activity of dopamine neurons and are the primary brake on dopaminergic activity (Barrot et al., 2012). GABAergic neurons intermingled amongst dopamine neurons in the VTA project to the NAc and LHb, as well as sending local collaterals (Brown *et al*., 2012; Root *et al*., 2014; Taylor *et al*., 2014). Thus, it is likely that in previous studies examining the influence of VTA GABAergic on disinhibition of dopamine neurons, that they are in fact stimulating tVTA/RMTg neurons located caudally to the VTA (van Zessen et al., 2012; Nieh et al., 2015, 2016).

Female mice show decreased asIPSC amplitude but not frequency of asychronous events from the LH GABA – tVTA/RMTg GABA projection compared to male mice, consistent with a decrease in number, function, or subunit composition of postsynaptic GABA_A_ receptors at LH GABA-tVTA/RMTg synapses. Notably, these effects were only observed during the asynchronous period 350 ms post optical stimulation, but not later, suggesting that this effect is specific to LH GABAergic inputs. LH GABA synapses were depressing at tVTA/RMTg neurons, there was a similar paired pulse ratio across all pulses. These results indicate that short term dynamics are similar at male and female tVTA/RMTg neurons with robust depression in both sexes.

Activation of LH GABAergic terminals decreases firing of tVTA/RMTg GABAergic neurons in both male and female rats. This effect was most pronounced within the first 5 pulses of the 20 Hz stimulation and largely driven by increased latency to fire rather than a decrease in AHP. Typically, a low threshold and fast inactivating K+ current (*I*_A_) underlies the latency to fire as it is activated during hyperpolarization and opposes membrane depolarization resulting in a delay to reach firing threshold (Kanold & Manis, 1999). Thus, optical stimulation of the LH GABAergic input likely hyperpolarizes tVTA/RMTg neurons leading to activation of *I*_A_ and a subsequent increased latency to fire (Kanold & Manis, 2005). One limitation with this experiment is that it is possible that ChR2 is variably expressed in different mice that could lead to potential group differences. We have tried to mitigate this by showing the relative eYFP reporter fluorescence is not different between group. However, this technique does not distinguish between eYFP fluorescence in terminals that synapse onto the recorded neuron and those that do not. Notably, we did not observe a difference in proportion of recorded tVTA/RMTg GABAergic neurons that did not produce an oIPSC in response to optical stimulation of LH GABAergic terminals between groups. Furthermore, there were no significant differences in paired pulse ratio in male vs female controls suggesting that differential ChR2 expression is not altering release properties in control mice.

### Effects of fasting on LH GABA – tVTA/RMTg GABA synapses

To examine the effects of optical stimulation of LH GABA inputs to the tVTA/RMTg, we recorded optically evoked IPSCs in ACSF where extracellular calcium was replaced with strontium which desynchronizes vesicular release to allow for the resolution of quantal synaptic events (Bekkers & Clements, 1999). The resultant local input specific asIPSC amplitude provides a measure of postsynaptic efficacy, whereas asIPSC frequency represents the number of connections or presynaptic function (Bekkers & Clements, 1999; McGarry & Carter, 2017). An acute overnight fast decreased the amplitude of LH GABA-mediated asIPSCs onto tVTA/RMTg neurons of male and female mice and decreased the frequency of asIPSCs of tVTA/RMTg neurons of female mice. This effect occurred within the first 350 ms after stimulation during the asynchronous release period. Thus, the local inhibitory drive onto tVTA/RMTg neurons after LH GABA stimulation is decreased with fasting, likely due to decreased synaptic strength and/or a possible decrease in number of local connections. Consistent with this, we observed a decrease in the optically evoked amplitude, the RRP size, and the steady-state oIPSC amplitude, of the LH GABAergic input to tVTA/RMTg GABA neurons after fasting in male and female mice that could contribute to the decrease in synaptic strength after fasting. Notably, unlike decreased frequency of asIPSCs in fasted female mice, fasting did not alter mIPSC frequency in male or female mice. This discrepancy could be due to the LH GABAergic inputs to tVTA/RMTg neurons provide only a smaller proportion of total inhibitory inputs to the tVTA/RMTg neurons, and thus a change is not detected when examining the contribution of all inhibitory synapses on mIPSC, or alternatively, different inhibitory inputs may have opposing effects. For example, while the LH GABAergic input is decreased with fasting, other inhibitory inputs may be strengthened with fasting resulting in no change detected when examining the fasting effect on all inhibitory synapses. Regardless, fasting appears to selectively reduce LH GABA release onto tVTA/RMTg neurons in female mice.

The reduction of firing of tVTA/RMTg neurons induced by optical stimulation of LH GABA inputs is diminished after an acute fast. However, approximately half the population of tVTA/RMTg neurons experienced depolarization block in response to a depolarizing step, yet in fasted mice, the number of cells with depolarization block was reduced by optical stimulation. This suggests that fasting may preserve the function of the LH GABA to tVTA/RMTg GABA input under depolarized conditions. The LH GABA input inhibits firing of tVTA/RMTg GABA neurons, but when the cell becomes depolarized for long periods of time leading to depolarization block, the LH input counters this and preserves firing and would allow for continued fine control of the circuit. Another possibility is that due to different intrinsic properties of tVTA/RMTg GABAergic neurons, some neuronal subtypes may be more susceptible to depolarization block in addition to possible fasting effects on intrinsic membrane properties. Although we did not observe an effect of fasting on depolarization block, the ability of optical stimulation of LH GABAergic terminals to reduce depolarization block in fasted mice may be due to an effect of fasting on intrinsic membrane properties in some cellular subtypes in addition to modulation of presynaptic terminals. A third possibility is that fasting-induced changes to the steady-state oIPSC may involve activation of extrasynaptic δ-subunit containing GABA_A_ receptors that underlie tonic GABA currents (Belelli *et al*., 2009). Tonic GABA currents play a role in modulating the excitability of neurons (Semyanov *et al*., 2004), and this may be one mechanism by which fasting can reduce the excitability of tVTA/RMTg GABA neurons. Future studies should address if fasting changes the tonic GABA inhibition of tVTA/RMTg GABAergic neurons of male of female mice.

Given that fasting increases the motivation to approach and consume food (Burnett *et al*., 2016; Godfrey & Borgland, 2020), it may appear counterintuitive that fasting does not change, but preserves the functional output of the LH GABA-tVTA/RMTg GABA synapses. While activation of LH GABAergic neurons produced consummatory behaviour in most (Nieh *et al*., 2015; Navarro *et al*., 2016; Marino *et al*., 2020), but not all paradigms (Burnett et al., 2016), its projections to the VTA likely drive behavioural activation. Stimulation of LH_GABA_-VTA inputs disinhibit VTA dopamine neurons and increases dopamine release in the NAc, leading to a promotion of motivated behaviours, including feeding, social interaction, generalized approach behaviours (Nieh et al., 2016) and reward predictions (Sharpe *et al*., 2017). Here, we demonstrated that fasting supresses firing of tVTA/RMTg GABAergic neurons, an effect on its own that would decrease inhibition of dopamine neurons. While in control mice, activation of LH GABAergic inputs reduced firing of tVTA/RMTg neurons, this effect was diminished during fasting due to a synaptic weakening of LH GABAergic inputs. Activation of LH GABAergic neurons in the unfasted state would typically disinhibit dopamine neurons (Nieh *et al*., 2015, 2016; Barbano *et al*., 2016). However, during fasting, even though LH GABAergic input to tVTA/RMTg neurons is weakened, fasting suppresses firing of tVTA neurons, and these effects together might still lead to a disinhibition of dopamine neurons that could promote food intake.

Notably, in our study, as well as those of others (Nieh *et al*., 2015, 2016; Barbano *et al*., 2016), some ChR2-eYFP expression was observed in the zona incerta (ZI). ZI neurons make monosynaptic projections to VTA dopamine neurons (Ogawa *et al*., 2014). Activation of ZI projections to the VTA can increase initiation of food intake (de Git *et al*., 2021), and thus it is possible that this input could be regulated by energy availability. However, it is unknown whether ZI neurons project to GABAergic neurons within the VTA or to those of the tVTA/RMTg. Future studies should address if ZI neurons indirectly modulate VTA dopamine neurons in different energy states.

In summary, we recorded from the caudomedial region of the VTA, a region containing almost exclusively GABAergic neurons, which are representative of the tVTA/RMTg population. Under sated conditions, activation of the LH GABA input suppresses firing of tVTA/RMTg GABA neurons. However, during fasting, the LH GABAergic input to VTA neurons is weakened, minimizing the hyperpolarizing effect of this input on tVTA/RMTg GABAergic neurons, such that firing of tVTA/RMTg is not changed to the same extent. Thus, the functional output of LH GABA neurons to disinhibit dopamine neurons is likely preserved in fasting leading to behavioural activation. An alternative hypothesis is that suppressed firing activity of tVTA/RMTg GABA neurons during fasting could be a mechanism associated with brain energy conservation during food scarcity, as has been observed in the neocortex which allowed for the significant conservation of ATP (Padamsey *et al*., 2021). However, as in Padamsey *et al*., 2021, synaptic functioning is preserved and activation of LH GABAergic inputs to tVTA/RMTg GABAergic neurons can override this fasting-induced suppression of synaptic strength. Future studies should aim to address either of these hypotheses.

## Additional Information

### Data Availability

Data is available upon request to the corresponding author.

### Author Contributions

NG performed electrophysiological, immunohistochemical and fasting experiments. MQ performed immunohistochemistry experiments. NG and SLB wrote first drafts of the manuscript. NG and SLB revised the manuscript. All authors approved the final version of the manuscript and are accountable for all aspects of the work.

## Acknowledgements

The authors would like to acknowledge the Hotchkiss Brain Institute advanced microscopy facility for their technical support.

### Land acknowledgement

This work was conceived and performed at The University of Calgary, located on the traditional territories of the people of the Treaty 7 region in Southern Alberta, which includes the Blackfoot Confederacy (including the Siksika, Piikuni, Kainai First Nations), the Tsuut’ina, and the Stoney Nakoda (including the Chiniki, Bearspaw, and Wesley First Nations). The City of Calgary is also home to Metis Nation of Alberta, Region 3.

